# A high-throughput RNA-seq approach to profile transcriptional responses

**DOI:** 10.1101/018416

**Authors:** G.A. Moyerbrailean, G.O. Davis, C. Harvey, D. Watza, X. Wen, R. Pique-Regi, F. Luca

## Abstract

In recent years, different technologies have been used to measure genome-wide gene expression levels across many types of tissues and in response to *in vitro* treatments. However, a full understanding of gene regulation in any given cellular and environmental context combination is still missing. This is partly because analyzing tissue/environment-specific gene expression generally implies screening a large number of cellular conditions and samples, without prior knowledge of which conditions are most informative (e.g. some cell types may not respond to certain treatments). To circumvent these challenges, we have established a new two-step high-throughput and cost-effective RNA-seq approach: the first step consists of gene expression screening of a large number of conditions, while the second step focuses on deep sequencing of the most relevant conditions (e.g. largest number of differentially expressed genes). This study design allows for a fast and economical screen in step one, with a more profitable allocation of resources for the deep sequencing of re-pooled libraries in step two. We have applied this approach to study the response to 26 treatments in three lymphoblastoid cell lines and we show that it is applicable for other high-throughput transcriptome profiling requiring iterative refinement or screening.

## Introduction

Recent studies have shown that the environment has played a major role in shaping the current distribution of allele frequencies in human populations.^1–3^ For example, one of the most striking cases of the clinical impact of human adaptations to dietary changes is the lactase persistence phenotype.^4^ These studies have investigated adaptations to the macroscopic (e.g. climate, resource availability, pathogen exposure) environment in humans. However, to learn about molecular mechanisms underlying the organismal response to environmental changes, it is necessary to focus at the cellular level. The cellular environment is determined by the presence of other cell types (e.g. in non-homogeneous tissues) and by the complex of stimuli (e.g. hormonal and metabolic) that a cell is exposed to. In this study, we define a specific environment as an agent that is introduced in the culture medium, can potentially change the state of the cell and is measurable at the molecular level. Examples include, agents secreted by nearby cells, hormones and metabolites secreted by other organs, pollutants, drugs or micronutrients absorbed by the organisms. These cellular environments are a complex function of organismal-level environmental exposures, which should have a more tractable influence on sub-cellular phenotypes such as gene expression. For example, environmental stress (physical or emotional) changes blood glucocorticoids level (a steroid hormone), which in the cell induces major changes in global gene-expression patterns mediated by the activation of the glucocorticoids receptor (GR).^5, 6^ Despite few additional examples, the molecular mechanisms underlying regulation of gene expression in different environmental conditions are still poorly understood. Functional genomics data collected by large consortia efforts, such as ENCODE (http://genome.ucsc.edu/ENCODE/), the Roadmap Epigenome (http://www.roadmapepigenomics.org/) have provided large amounts of information on tissue specific regulatory regions of the human genome. However, we are still missing a full understanding of the elements that regulate gene expression in any given cellular and environmental context.

The first step in understanding the regulatory mechanisms underlying a cell’s response to environmental perturbations is a comprehensive characterization of the transcriptional response across several environments (treatments and cell types). Current experimental setups are costly and laborious and have allowed only focusing on the analysis of a particular cellular environment in any given experiment. The most notable exception is the Connectivity Map initiative,^7^ which characterized the transcriptional response to 164 small-molecule perturbagens in four cancer cell lines. Here we have developed an approach that allows a similar throughput, and uses RNA-seq rather than microarrays, with specific advantages: for example, with RNA-seq it is possible to investigate a wider dynamic range of expression levels, with digital sensitivity; it is also possible to study transcript isoforms and RNA species that are not represented on common arrays, such as LincRNAs. Recently, Narayan and colleagues^8^ have developed a high throughput (META) RNA profiling method, that allows for parallel analysis and multiplexing of a large number of samples. However, the tag and PCR-based method in the current protocol, only allows for the quantification of known RNA species and prevents some RNA-seq specific applications such as isoforms quantification and allele-specific analysis. Since the development of RNA-seq technology,^9–11^ a variety of protocols have been introduced to investigate specific biological problems, for example, direct RNA sequencing,^12^ allows sequencing of RNA molecules skipping cDNA synthesis and can thus analyze short, degraded and/or small quantity RNA samples; another example of fast and automatized RNA-seq protocols is the Tn-RNA-seq^13^ approach, which uses transposase-based incorporation of sequencing adaptors in cDNA libraries.

Independent of the RNA-seq technology used, here we present a new cost-effective two-step strategy that uses the ability to index and pool many (96 or more) RNA-seq libraries in parallel. This strategy allows the researcher to rapidly screen a large number of sample conditions and strategically allocate sequencing resources to in depth analysis only for the relevant cases. We demonstrate this approach by exploring the transcriptional response to a wide panel of environmental perturbations (26 treatments) in three LCL samples. The results show that our approach should also be applicable to similar scenarios requiring high throughput screening across multiple cell lines, treatments, time points and/or patient samples in a variety of contexts, such as: population genetic studies, pharmacological drugs testing and cancer transcriptome profiling.

## Results

### The two-step approach

We have developed a new high-throughput two-step approach to characterize the transcriptional response to environmental perturbations through RNA-seq. An outline of the approach is presented in Figure 1. In step one we characterize global changes in gene expression. Here we used a modified RNA-seq protocol (see Methods) better suited for our specific application, but similar results can be achieved with popular commercial RNA-seq kits that allow for high multiplexing (96-well plate format) such as the Illumina TruSeq Stranded mRNA HT Sample preparation kit or the NEBNext Ultradirectional (NEB) library preparation kit. Many of these commercially available kits can work with liquid handling robots that automatize the majority of the experimental steps (e.g., Beckman Coulter Biomek FXp, Eppendorf epMotion 5075, and others).

**Figure 1.**
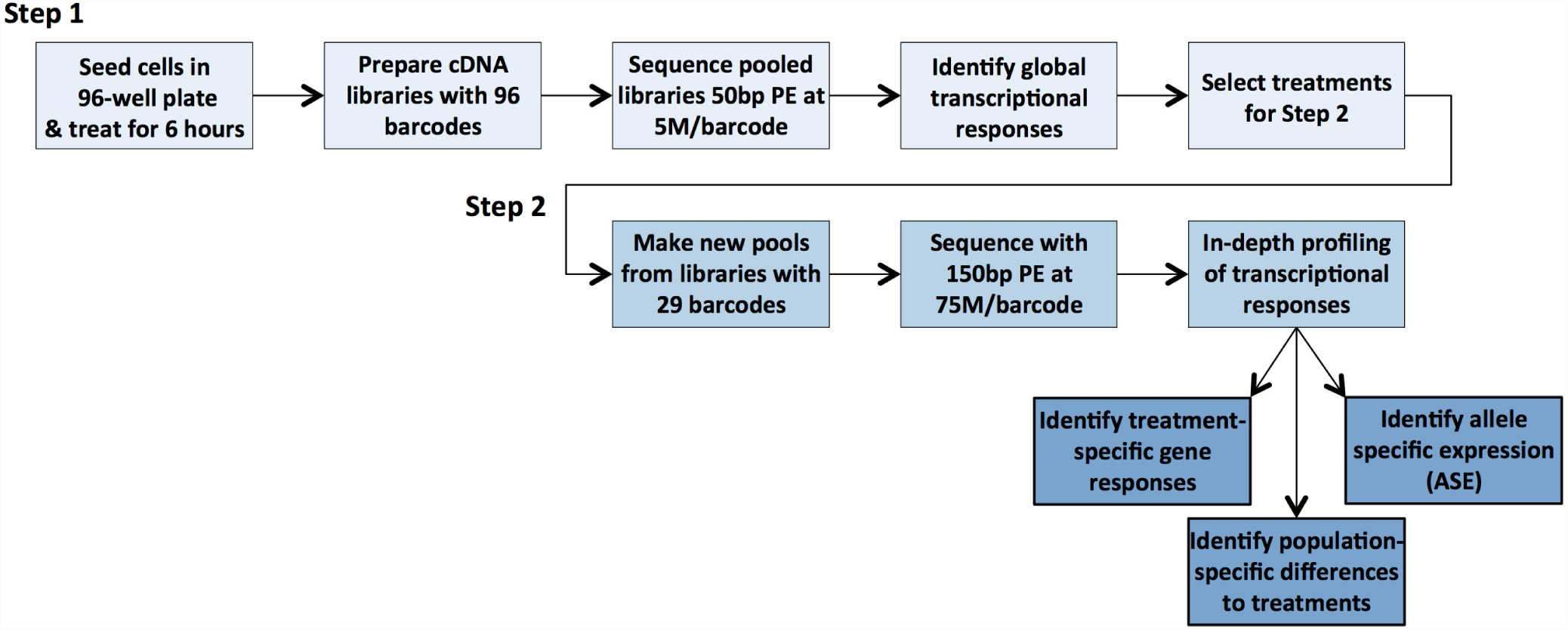
Workflow of the two-step approach.

In the first step, all samples are experimentally processed in parallel, from tissue culture and treatments to library preparation, thus minimizing variation from testing hundreds of conditions at the same time. Additionally, high multiplexing allows reducing the number of controls that would need to be repeated across different treatment batches in a less multiplexed experimental set up (e.g. 93 treatments plus 3 controls). A 96-libraries pooling and shallow sequencing strategy is then used to minimize the amount of resources used in the screening step. Most RNA-sequencing studies that only require gene expression quantification are currently collecting tens of millions of reads per sample. A recent report by Hou *et al*^14^ shows that gene expression for high abundant transcripts can be reliably quantified with less than five million reads. Here we demonstrate that similar sequencing depth also allows detecting global and biologically relevant gene expression changes and can be used to identify relevant conditions to follow up in step two. Furthermore, even for study designs that require deep sequencing of large number of samples (e.g. 96), our two-step approach allows using the first step to QC the libraries, before investing in deep sequencing efforts.

For the second step, we do not need to prepare new libraries, and we can simply repool a selection of the initial libraries, without additional experimental costs. Furthermore, we can optimize library concentrations to pool in order to achieve even representation of individual libraries. This is done by calculating a digital library concentration from the sequencing run performed in step one. Note that this digital library concentration is the fraction of reads from the total sequenced in the pool, and it naturally takes into account potential differences across the libraries in sequencing output (e.g., due to Flow cell cluster formation in Illumina sequencing machines) (see Methods, Equation 1). Even in situations where deep sequencing data are to be collected for all samples, using a two-step approach allows for optimizing pooling ratios and efficient allocation of sequencing resources across samples. The greater sequencing depth in step two is carried out by capitalizing on the resource savings made possible by the shallow sequencing scheme applied in step one.

Below we present an application of the two-step approach to analyze the response to 26 environmental perturbations in LCLs.

#### Step one: Identifying global changes in gene expression from low-coverage data

To characterize the response to treatments, cells were treated, while in mid-log phase exponential growth, with the panel of treatments listed in Table 1, for 6 hours. Cells from all treatment conditions, including the vehicle controls, were cultured and harvested in parallel at the same time point. This is different from studies that collect genomic data prior and post treatment (with the treatment being done in vivo or in vitro) and allows for a better control of technical noise, or biological variation that is independent of the treatment. For example, we were able to account for temporal changes in gene regulation that are independent of the treatment (e.g. changes in cell cycle phase over time, reagent batch effect). Furthermore, to achieve greater confidence and accuracy to measure baseline gene expression, for each LCL sample, the control treatments were performed in triplicates. For all stages of sample preparation we have used a 96-well plate study design (3 samples and 32 treatment conditions): from cell culturing to RNA extraction and library preparation, thus facilitating increased sample processing throughput.

**Table 1.** Treatments used in step one.

| Treatment | Common Name | Control | Concentration<sup>†</sup> |
| --- | --- | --- | --- |
| Ascorbic acid | Vitamin C | Media | $1.00 \times 10^{-5}$ |
| Biotin | Biotin | Media | $4.75 \times 10^{-10}$ |
| Nicotinic Acid | Vitamin B3 | Media | $1.50 \times 10^{-5}$ |
| Pantothenic Acid | Vitamin B5 | Media | $1.00 \times 10^{-7}$ |
| Pyridoxine | Vitamin B6 | Media | $1.00 \times 10^{-5}$ |
| Retinoic Acid | Vitamin A | Ethanol | $1.00 \times 10^{-8}$ |
| Tocopherol | Vitamin E | Ethanol | $5.00 \times 10^{-5}$ |
| Plumbagin | Vitamin K3 | Ethanol | $1.00 \times 10^{-6}$ |
| Aldosterone | Aldosterone | Ethanol | $1.00 \times 10^{-5}$ |
| Progesterone | Progesterone (C1) | DMSO | $1.00 \times 10^{-6}$ |
| Progesterone | Progesterone (C2) | Ethanol | $1.00 \times 10^{-5}$ |
| Beta-Estradiol | Estrogen | Ethanol | $1.00 \times 10^{-5}$ |
| Dexamethasone | Dexamethasone | Ethanol | $1.00 \times 10^{-5}$ |
| Caffeine | Caffeine | Media | $1.16 \times 10^{-3}$ |
| Nicotine | Nicotine | Media | $6.16 \times 10^{-4}$ |
| Copper (II) Chloride | Copper | Media | $6.00 \times 10^{-5}$ |
| Iron (III) Chloride | Iron | Media | $5.00 \times 10^{-3}$ |
| Manganese (II) Chloride | Manganese | Media | $3.00 \times 10^{-3}$ |
| Molybdenum (V) Chloride | Molybdenum | Media | $5.00 \times 10^{-4}$ |
| Sodium Selenite | Selenium | Media | $1.00 \times 10^{-5}$ |
| Zinc Chloride | Zinc | Media | $8.00 \times 10^{-5}$ |
| Tunicamycin | Tunicamycin | DMSO | 2 μg/mL |
| PM 2.5 (Detroit) | PM 2.5 | Media | 5 μg/mL |

<sup>†</sup> All concentrations are in molarity (M) unless otherwise specified.

To identify DE genes we used the method implemented in the software DEseq2, which estimates variance-mean dependence in the read counts for each gene and tests for differential expression based on a model using the negative binomial distribution.

Each treatment was matched to the appropriate vehicle control (Table 1) for this analysis. However, when comparing pairs of controls to each other we did not detect any DE genes (10% BH-FDR, Figure S1). To assess the calibration of the tests for differential expression on low coverage data, we used QQ-plots and observed that in most cases the tests are well calibrated (Figure 2 and Figure S2).

**Figure 2.**
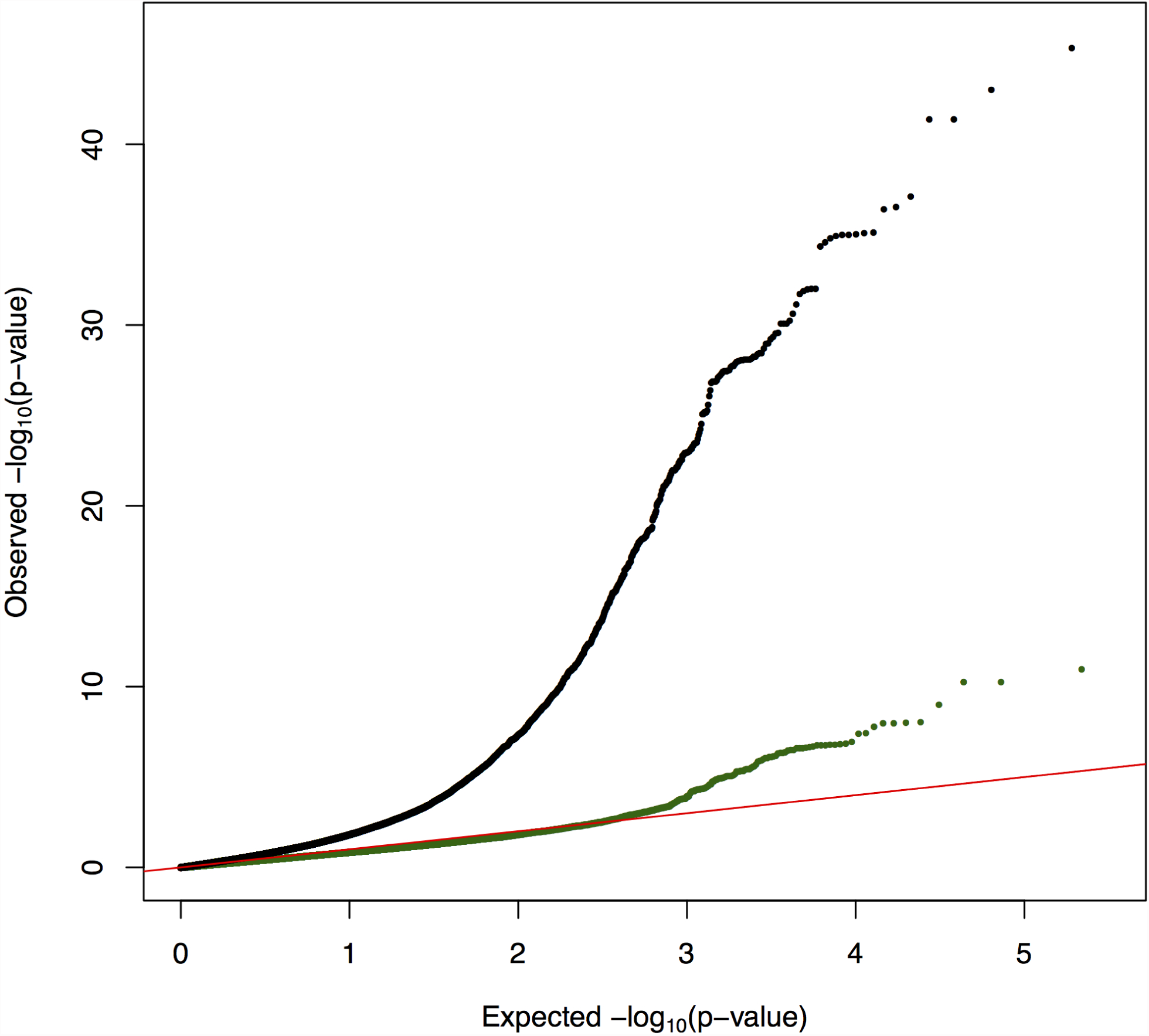
QQplot of the *p*-value distribution for DE genes in response to dexamethasone (black) and estrogen (green). Additional QQ plots from step one and step two are available in the supplements.

We next asked whether our ability to detect DE genes may depend on sequencing depth. Figure 3 shows that the number of DE genes is not correlated with sequencing depth across three individual samples for each treatment. For example, we detect *<*100 DE genes in response to vitamin B6 and vitamin E, even though these are the treatments for which we collected the largest number of reads. On the other hand, we identified thousands of DE genes for iron and tunicamycin, which are among the treatments with less coverage.

**Figure 3.**
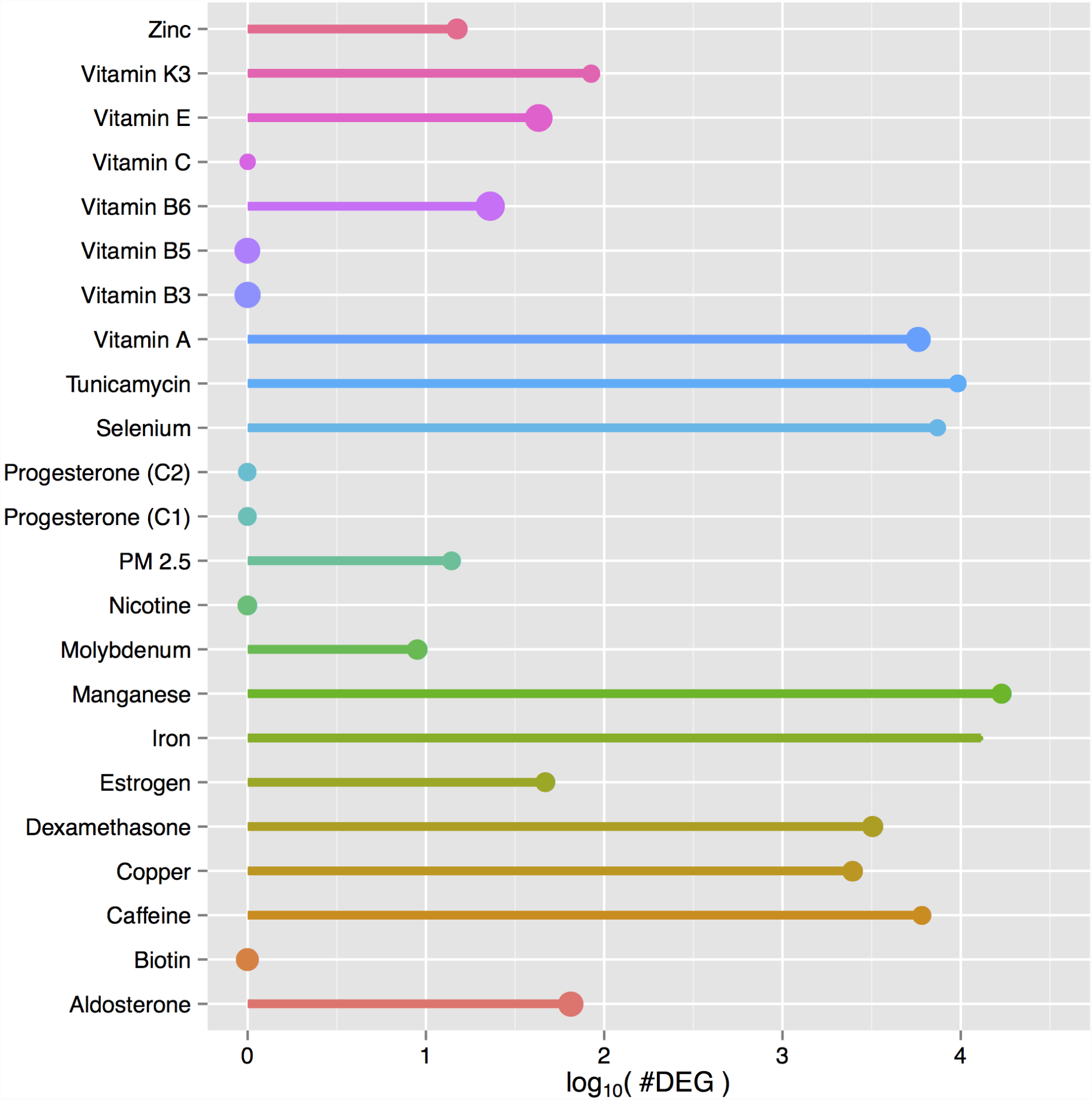
DE genes and sequencing depth. Bubble plot of DE genes (10% BH-FDR). The size of the bubble indicates the total number of reads after filtering across treatment samples.

We selected the panel of treatments to include dexamethasone and estrogen as prior experiments performed by our group (see Luca *et al*, 2013,^15^ Maranville *et al*, 2011^5^) showed a strong and null transcriptional response to these compounds, respectively. After running DESeq2, we identified 3212 DE genes in response to dexamethasone, while only 47 DE genes were detected in cells treated with estrogen, thus confirming previous results (Figure 2).

To identify major similarities and differences in the transcriptional response to our panel of treatments, we performed hierarchical clustering on the transcript expression data for each treatment (expressed in FPKMs). Figure 4 shows a heatmap of the correlation matrix across all treatment conditions and samples. Some key features appear evident even with low sequencing depth: control samples cluster together; treatments that induce a strong response are distinct from all other treatments and controls, and show a clear pattern where the three samples for each treatment condition cluster very tightly.

**Figure 4.**
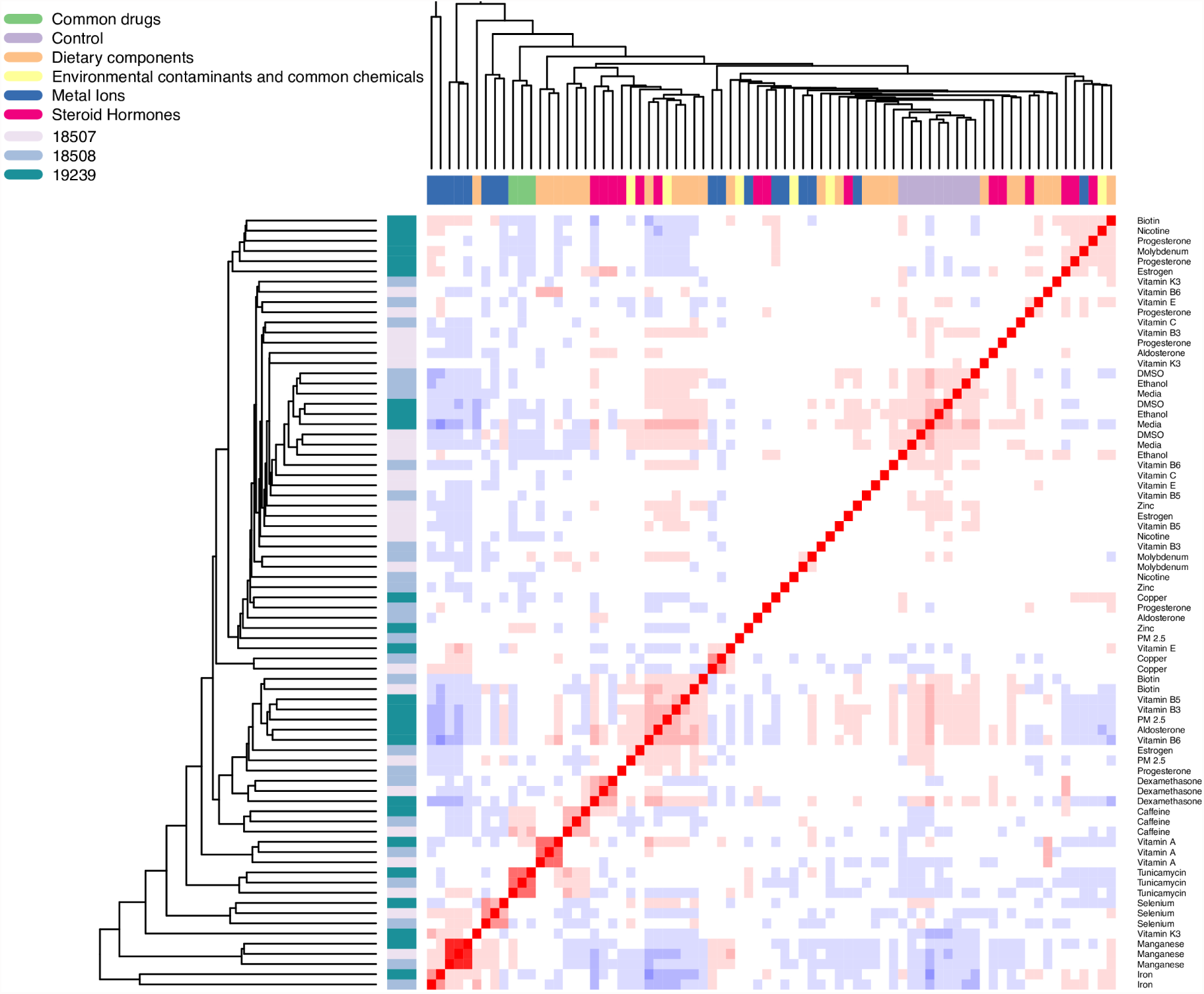
Heatmap and hierarchical clustering of gene expression levels. Gene expression levels (FPKMs) were clustered for each sample (row coloring) and treatment (column coloring, row labeling) combination. The dendrogram shows the Euclidian distance between samples, while the heatmap shows the Pearson correlation (red = 1, blue = *−*1).

Given the high number of DE genes observed for certain treatments, we asked whether they could indicate that the cell is undergoing a cytotoxic response. To this end we compared the transcriptional response from RNA-seq data to the cytotoxic response measured in viability assays. To measure cell viability we used the Promega Glo-Max assay, and compared ATP production measured in relative luminescence units in treatment and control cells. We observed a significant negative correlation between number of DE genes and change in cell viability after 48 hrs treatment (Spearman *ρ* = −0.48, *p* = 0.02). This suggests that when an extremely large number of DE genes is observed, it is indicative of major changes in the cell physiological state, which ultimately may lead to cell death. For example, the largest number of DE genes was observed for treatments such as iron (12979) and manganese (16990), which were administered at supra-physiological doses (Figure S2).

Overall the results from step one show that even from low sequencing depth data it is possible to identify biologically meaningful global changes in gene expression that are relevant to assess the cellular response to environmental perturbations.

#### Step two: Analysis of gene regulation in response to environmental perturbations

The information collected in step one of our approach can be most effectively used to re-pool individual libraries by selecting the treatment conditions biologically relevant for the system under-study. As a proof of principle, we selected four treatment conditions (vitamin A, copper, selenium, iron) for deeper sequencing (75M reads/sample) in all three cell lines, to investigate the transcriptional response to these environmental perturbations with greater resolution.

One of the challenges when sequencing highly multiplexed pools of libraries is achieving even coverage across samples. Figure 5 shows density plots of sequencing depth across shallow and deep sequencing samples. The distribution of sequencing depth is a function of factors related to the sequencing technique and instrument, including, for example, efficiency in cluster generation on an Illumina sequencer. It is possible to account for these factors when determining pooling concentrations for the deep sequencing pool. We developed a formula (Equation 1) that uses information from the low coverage data to learn about “read” concentration per library and also accounts for the sequencing output of each individual sequencing run (see Methods). This is much better than any standard library quantification approach, because we have a “digital” count of the actual reads that contribute to generate clusters on the flow-cell per unit of volume of the library. As expected, in the 30 pooled libraries from step two, we observe a much tighter distribution of sequencing depth.

**Figure 5.**
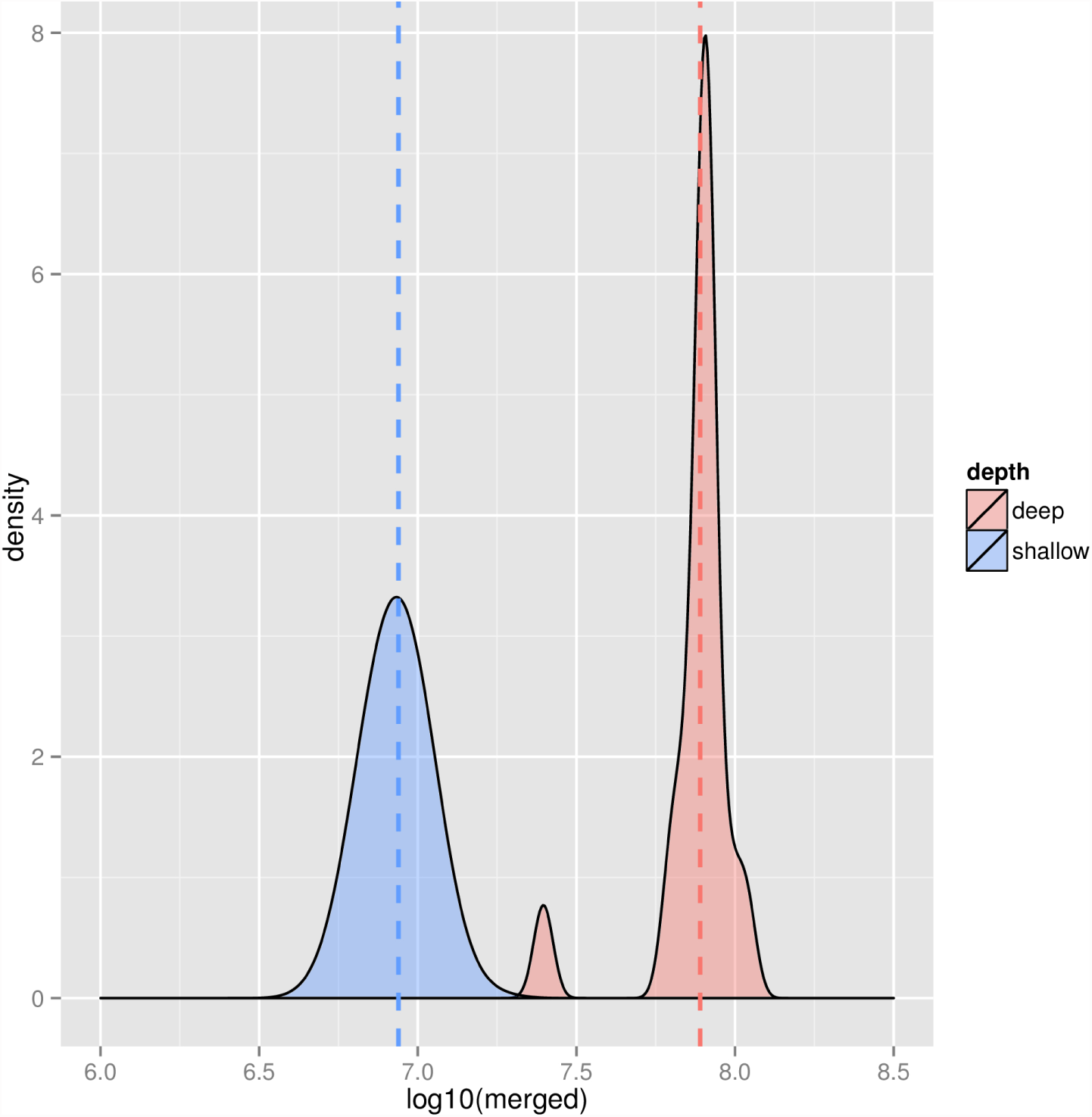
Density plot of raw (unfiltered) sequencing depth across individual barcoded samples for the shallow sequencing (blue) and across treatments for the deep sequencing (red) runs. Note that in step two, to achieve even representation of sequencing depth across treatment conditions, each replicate of the control samples was pooled at one third of the other treatments. Dotted lines indicate mean sequencing depth.

We then used DEseq2 to identify DE genes in the deep sequenced libraries. Table 2 shows the number of DE genes, and their direction of expression change. We found that transcript fold change is highly correlated between the shallow and deep sequencing experiments for the same treatment (Spearman *ρ >* 0.7, Figure 6), which confirms that gene expression changes detected from shallow sequencing can be used to identify biologically relevant treatments for follow up studies. As expected, with deep sequencing data we can identify transcriptional changes at greater resolution. Figure 7 shows the increase in number of DE genes as a function of sequencing depth in step one and step two.

**Table 2.**
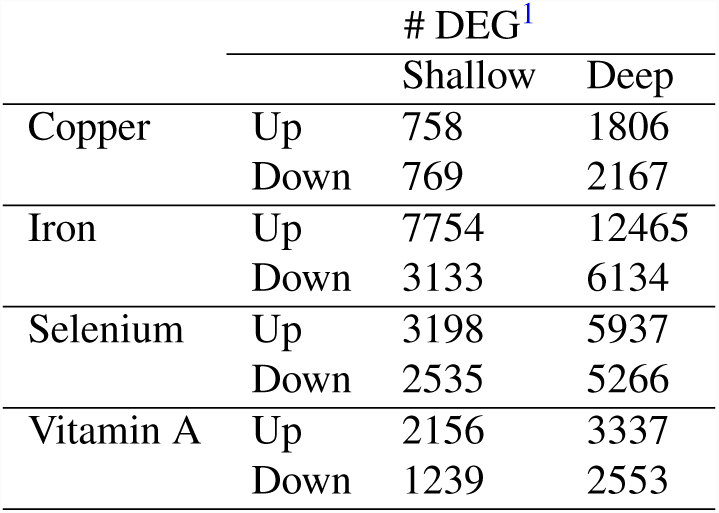
Differentially expressed genes identified in step two.

**Figure 6.**
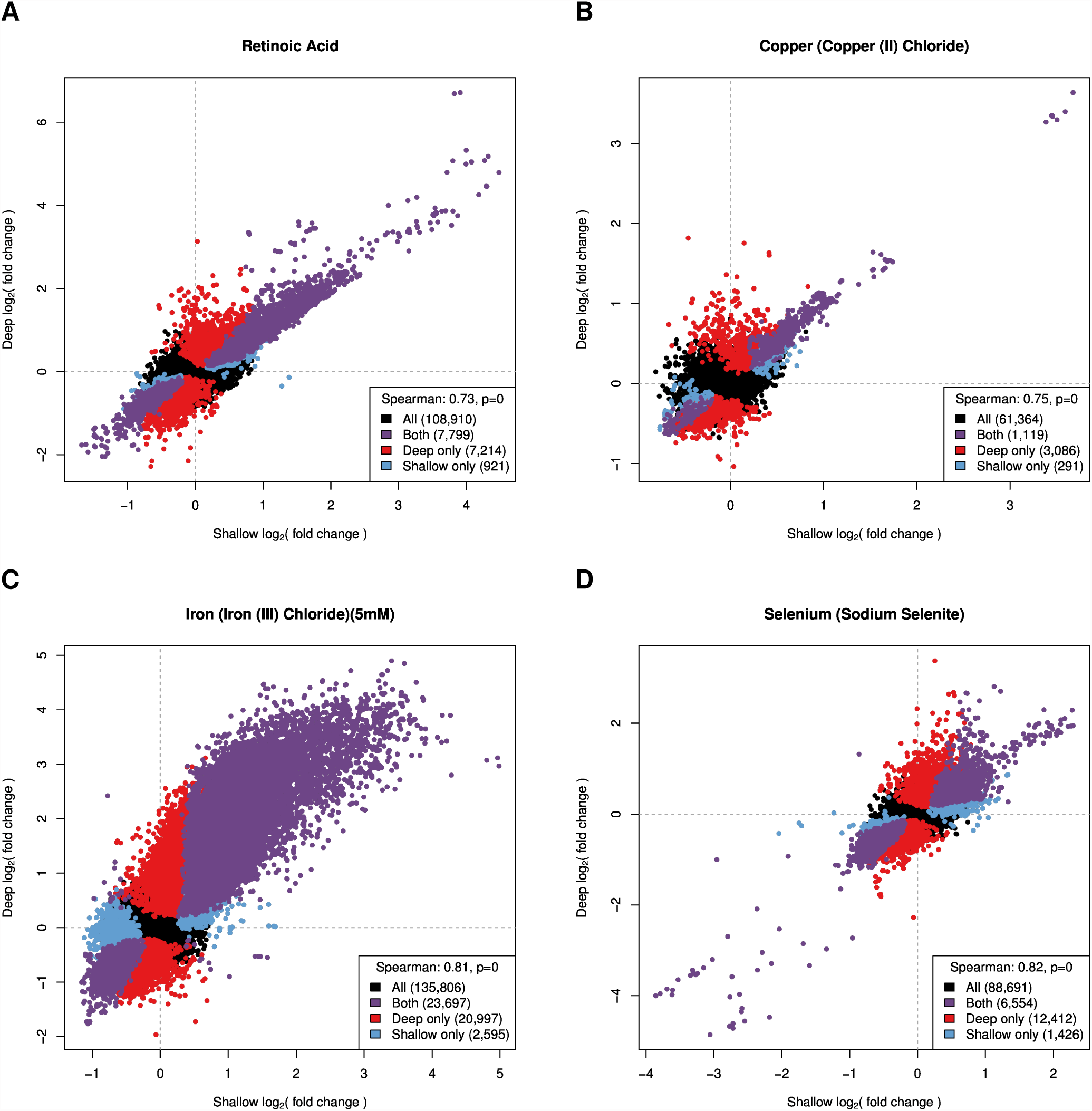
Correlation in the transcriptional response between shallow and deep sequencing. Plotted is the log_2_(fold change) for each transcript calculated from shallow and deep sequencing data for the four treatments analyzed in step two. Colored points represent transcripts differentially expressed at 1% BH-FDR. Vitamin A (A), copper (B), iron (C), selenium (D).

**Figure 7.**
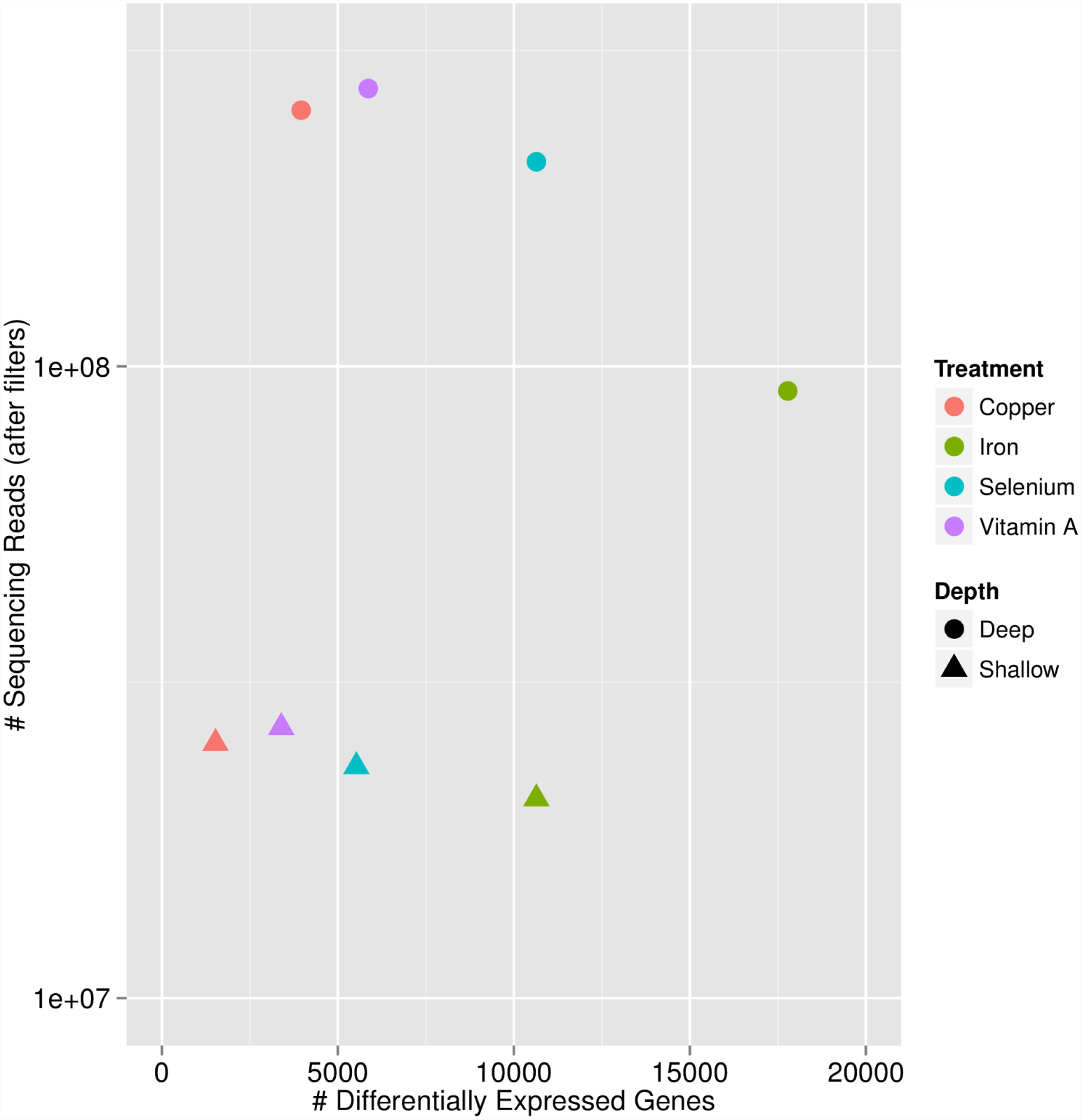
Comparison of sequencing depth and number of differentially expressed genes (10% BH-FDR) between shallow and deep sequencing runs.

To investigate similarities in the transcriptional response to these four treatments, we calculated pairwise Spearman rank correlations on the transcript fold change. We observed that responses to metal ions (copper, selenium, iron) tend to be more highly correlated with each other, compared to vitamin A. The highest correlation was observed between copper and iron (0.43, *p* = 0). This suggests that LCLs respond to these treatments through similar gene regulatory pathways.

To further investigate the regulatory pathways altered during the response to these treatments, we performed GO enrichment analysis using the DAVID online tool^16^ and focusing on biological processes (5% BH-FDR, Supplementary Tables S1-S8). We observed that upregulated genes in response to vitamin A are enriched for the immune response and related processes (e.g. leukocytes and lymphocytes activation), which is in line with the known role of vitamin A as an activator of immune function.^17^ Upregulated genes in response to copper are enriched for genes involved in the protein ubiquitination biological processes, and the same result is observed for upregulated genes in response to selenium. This supports the observation that these two metal ions elicit very similar transcriptional responses, which are clearly distinct from the response induced by treatment with vitamin A. However GO enrichment analysis also points to an anti-inflammatory role for selenium, as down-regulated genes in response to this metal ion are enriched for leukocytes activation. Finally, genes upregulated in response to iron are enriched for metal ion transport and cell-cell adhesion among the top biological processes, while down-regulated genes are enriched for RNA and DNA metabolic processes as well as key cellular processes such as mitosis. These last enrichments reflect the observed cytotoxicity of the iron treatment we performed on the cells.

## Discussion

We have developed a novel high-throughput and cost-effective approach to screen and analyze the transcriptional response to a large number of environmental perturbations through RNA-seq. This approach consists of two steps, where only the first step requires cell culture experiments and library preparation and allows for a fast and economical screen of a large number of environmental conditions that are followed up in the second step through deep sequencing of re-pooled libraries.

We have shown that shallow sequencing of 96 pooled libraries allows identifying, with minimal costs (approx $60/sample, including library preparation and sequencing), the most interesting conditions while capturing biologically relevant and informative gene expression changes. This removes the burden of deep sequencing uninformative libraries in pilot studies. We have presented an application of this approach to analyzing 23 environmental perturbations and appropriate controls in three LCL samples. However, this approach can be successfully applied to other study designs where it is most economical to test a large number of conditions prior to further analysis of relevant ones. Examples of such applications include time-course experiments, where it may be relevant to initially screen a large number of timepoints, to identify the most relevant ones to follow up. At the population level, to identify population-specific responses, one could first test a large number of treatments in a few individuals in order to identify the treatments that elicit a strong response to justify treating large population samples. Finally, tissue specific profiling in disease patients, where step one would allow identifying the most differentially expressed tissues between patient and controls that justify deep sequencing to characterize specific altered pathways.

The second step of our approach can be designed to achieve varying levels of sequencing depth and read length, depending on the question being asked. Here we have used step two to validate the shallow sequencing step and learn about transcriptional changes in response to three metal ions and a vitamin/nuclear receptor ligand treatments.

Given the significant monetary savings allowed by step one, it is possible to invest in deep sequencing of step two pools to the degree necessary to answer specific biological questions. For example, using more cycles to get longer reads, may facilitate transcript isoforms detection and quantification.^18^ A sequencing depth of 80M reads or above combined with longer reads also helps in identifying allele specific expression (ASE) even in the absence of genotype information, as we recently showed in.^19^

With the availability of desktop sequencer instruments (such as the Illumina NextSeq500), this two-step protocol will allow for fast screening and in-depth analysis of relevant conditions in less than 1 week-time. Additionally, compared to microarray-based pilot studies with 96 samples, our approach allows for 40% savings (e.g. $9,600 with the least expensive microarray option vs $5,800), with subsequent optimal allocation of resources to meaningful biological conditions (in step two), thus reducing the amount of time and funds spent on unsuccessful pilot/exploratory studies.

## Methods

### Cell culture and treatments

Lymphoblastoid cell lines (LCLs) were purchased from Coriell Cell Repositories. Prior to the experiment, cells were cultured, at 37° and 5% CO2, in RPMI 1640 (Gibco), supplemented with 15% heat-inactivated fetal bovine serum and 0.1% Gentamycin. The following LCLs were used: GM19239, GM18507, and GM18508. LCLs were cultured in “starvation medium” composed of RPMI 1640, supplemented with 15% charcoal-stripped fetal bovine serum (CS-FBS) and 0.1% Gentamycin for four days. Cells were then treated with the treatment panel (Sigma Aldrich) in Table 1 for 6 hours.

### Sample Collection and mRNA isolation

Treated cells were collected by centrifugation at 2000 rpm and washed 2X using ice cold PBS. Collected pellets were lysed on the plate, using Lysis/Binding Buffer (Ambion), and frozen at -80°. Poly-adenylated mRNAs were subsequently isolated from thawed lysates using the Dynabeads mRNA Direct Kit (Ambion) and following the manufacturer instructions.

### A modified RNA-seq library preparation protocol

We modified the NEBNext Ultradirectional (NEB) library preparation protocol to use 96 Barcodes from BIOOScientific added by ligation, this allowed us to reduce the overall library preparation cost to $47/sample. Specifically, RNA-seq libraries were prepared using the NEBNext ultradirectional library preparation protocol, with the following changes. RNA was fragmented at 94° for 5 minutes to obtain fragments 200-1500bp in size. SPRI Select beads (Beckman Coulter) were used in all purification steps and size selection was performed to obtain 300-450bp fragments. After the cDNA synthesis, to the 65 *μ*L of dA-Tailed cDNA were added the following components: 15*μ*L of Blunt/TA Ligase Master Mix, 2.39*μ*L of BIOO Scientific Barcode Adaptors (1-96), 1.11 *μ*L of Nuclease-free water. The samples were incubated for 15 minutes at 20° in a thermal cycler. USER Excision and PCR Library Enrichment were performed according to the following protocol. To the size selected cDNA (20*μ*L) were added the following components: 3*μ*L of NEBNext USER Enzyme, 25*μ*L of NEBNext High-Fidelity PCR Master Mix, 2X, 2*μ*L of BIOO Scientific Universal Primer (12.5*μ*M). The individual libraries were quantified using the KAPA real-time PCR system, following the manufacturer instructions and using a custom-made series of standards obtained from serial dilutions of the phi-X DNA (Illumina). Pools of 96 samples from the first step were sequenced on two lanes of an Illumina HiSeq in fast mode to obtain 50bp PE reads, at the University of Chicago Genomics core. Alternatively, this could be run on one lane of the Illumina Next-Seq 500 (75 cycles, PE). Re-pooled libraries for step two were sequenced on 1 lane of the Illumina Next-Seq500 in High Output mode to obtain 150bp PE reads in the Luca laboratory.

### Calculating optimal re-pooling proportions

To calculate optimal re-pooling proportions after shallow sequencing, we first calculated the digital concentration of reads*/μ*L (*R*). For each sample *i* sequenced in step one, *R*_*i*_ is defined as the number of raw sequencing reads per *μ*L of pooled library. The re-pooling proportion for each sample *i* is then calculated using the following formula:

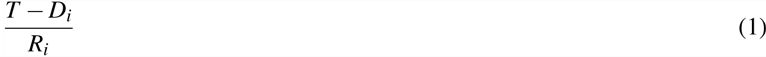

where *T* represents the total number of reads desired for each sample *i* (here 75M) and *D*_*i*_ represents the number of reads collected for sample *i* in previous runs. Changing the value for *D*_*i*_ and *R*_*i*_ also allows for iterative adjustments of pooling proportions in order to reach the desired total number of reads through multiple re-pooling and sequencing runs.

### RNA-seq data processing and differential gene expression analysis

Sequencing reads were aligned to the reference human genome hg19 using 

~~~
bwa mem
~~~

^20^ (http://bio-bwa.sourceforge.net). Reads with quality *<*10 and duplicate reads were removed using 

~~~
samtools rmdup
~~~

 (http://github.com/samtools/). We also removed two samples (barcodes) because the sequencing failed (extremely low number of reads, *<*50,000). Read counts covering each transcript were calculated using samtools and the Ensembl gene annotations. Counts data for transcripts with *>*20 reads were used to run DESeq2.^21^ To best account for overdispersion, the DESeq2 model was fit on all sequencing data simultaneously, rather than pairwise matching of treatments and controls. Each control-treatment pair was then matched from an experimental design matrix, and differentially expressed (DE) genes were determined as those with at least one transcript with a Benjamini-Hochberg controlled FDR^22^ (BH-FDR) of 10%. For step two, reads from multiple runs were merged after alignment (at the bam stage) and prior to applying any filter.

To perform hierarchical clustering of the expression levels across treatments, for each transcript in the Ensembl annotations, we calculated FPKMs from the number of reads covering the transcript. To control for potential confounders of expression data, a linear model was used to regress out effects from GC content, transcript length, and an interaction term between GC content and transcript length. These residuals were quantile normalized within each sample, and normalized within each individual by subtracting that individual’s average value per transcript across all treatments (10% trimmed mean).

### Viability Assays

To assess cell viability in response to the treatment panel, cells were exposed to each environmental stimulus and subsequently evaluated using the CellTiter-Glo Luminescent Assay (Promega Cat-G7570). LCLs were cultured and treated as described above, with the exception of being seeded into a 96-Well-Black tissue culture plate (Fisher). Treated plates were then incubated for 48 hours. After each incubation period, the CellTiter-Glo assay was performed according to the manufacturer protocol. The plate was then scanned in the Fluoroskan Ascent FL plate reader and luminescent signal acquired. For each treatment and control sample, at each time point, experiments were performed in triplicates on one LCL sample. Significant differential viability was assessed by a *t*-test comparing each treatment to the appropriate vehicle-control.

## Acknowledgements

We thank Athma Pai for helpful discussions at the early stages of the project, Stephen Krawetz for helpful comments on a preliminary version of the manuscript, the University of Chicago Functional Genomics Core for sharing the modified protocol for the KAPA real-time PCR system, the Wayne State University High Performance Computing Grid for computational resources and members of the Luca/Pique-Regi group for comments and advice. This work was supported by the National Institutes of Health [5R01GM109215 to F.L and R.P.]; and the American Heart Association [14SDG20450118 to F.L.].

## Author contributions statement

F.L. X.W. and R.P. conceived the experiments, G.D. and D.W. conducted the experiments, G.M. and C.H. analysed the results, F.L. and R.P. supervised data analysis, F.L. R.P. G.M. and G.D. wrote the manuscript. All authors reviewed the manuscript.

## Additional information

The authors declare no conflicts of interest.

